# Understanding the functional and structural differences across excitatory and inhibitory neurons

**DOI:** 10.1101/680439

**Authors:** Sun Minni, Li Ji-An, Theodore Moskovitz, Grace Lindsay, Kenneth Miller, Mario Dipoppa, Guangyu Robert Yang

## Abstract

One of the most fundamental organizational principles of the brain is the separation of excitatory (E) and inhibitory (I) neurons. In addition to their opposing effects on post-synaptic neurons, E and I cells tend to differ in their selectivity and connectivity. Although many such differences have been characterized experimentally, it is not clear why they exist in the first place. We studied this question in an artificial neural network equipped with multiple E and I cell types. We found that a deep convolutional recurrent network trained to perform an object classification task was able to capture salient distinctions between E and I neurons. We explored the necessary conditions for the network to develop distinct selectivity and connectivity across cell types. We found that neurons that project to higher-order areas will have greater stimulus selectivity, regardless of whether they are excitatory or not. Sparser connectivity is required for higher selectivity, but only when the recurrent connections are excitatory. These findings demonstrate that the differences observed across E and I neurons are not independent, and can be explained using a smaller number of factors.

---

Deep neural networks have become powerful tools to model the brain [Yamins and DiCarlo, 2016, Kriegeskorte, 2015]. They have been used to successfully model various aspects of the sensory [Yamins et al., 2014, Kell et al., 2018], cognitive [Mante et al., 2013, Wang et al., 2018, Yang et al., 2019], and motor system [Sussillo et al., 2015]. Deep networks have been particularly effective at capturing neural representations in higher-order visual areas that remain challenging to model with other methods [Yamins et al., 2014, Khaligh-Razavi and Kriegeskorte, 2014].

Demonstrating that these models can mimic the brain in some respects is only a starting point in the process of advancing neuroscience with deep learning. Once some relationship between artificial and biological networks is established, the model can be dissected to better understand *how* certain biological computations are mechanically implemented [Sussillo and Barak, 2013]. Although notoriously difficult to interpret, deep networks are still much more accessible than the brain itself. This is how deep networks can help address the “how” question. On top of that, deep networks can be used in a normative approach to answer the “why” question. Suppose we found that a certain architecture, objective function (dataset), and training algorithm could together lead to a neural network matching some features of the brain. Then we can ask why the brain evolved such features by testing which element of the architecture, dataset, and training is essential for such features to evolve in the artificial networks. For example, Lindsey et al. [2019] showed early layers of a convolutional neural network can recapitulate the center-surround receptive field observed in retina, but only when followed by an information bottleneck similar to the one that exists from retina to cortex. This finding demonstrates how realistic biological constraints can be used to explain the emergence of known properties of the brain.

Despite the many successes of applying deep networks to model the brain, standard architectures deviate from the brain in many important ways. Recently, neural networks that incorporate fundamental structures of the brain are becoming increasingly common [Kar et al., 2019, Miconi et al., 2018, Song et al., 2016]. One fundamental structural principle of the brain is the abundance of recurrent connections within any cortical area. Cortical neurons receive a substantial proportion of their inputs from other neurons in the same area [Harris and Shepherd, 2015]. Convolutional recurrent networks trained on object classification tasks can perform comparably to purely feedforward networks with similar numbers of parameters, while being able to better explain temporal responses in higher visual areas [Nayebi et al., 2018, Kar et al., 2019].

One of the most fundamental organizational principles of the cortex is the separation of excitatory (E) and inhibitory (I) neurons [Dale, 1935, Eccles et al., 1954]. Dale’s law states that each neuron expresses a fixed set of neurotransmitters, which results in either excitatory or inhibitory downstream effects. In addition to the difference in their immediate impacts on post-synaptic neurons, excitatory and inhibitory neurons differ in several other important ways. There are several times (4-10x) more excitatory neurons than inhibitory neurons [Hendry et al., 1987]. Neurons that project to other areas, the so-called “principal neurons”, are all excitatory in the cortex [Bear et al., 2007]. In the mammalian sensory cortex, excitatory neurons are overall more selective to stimuli than inhibitory neurons in the same area as extensively reported in mice [Kerlin et al., 2010, Znamenskiy et al., 2018] and to a lesser extent in other species [Wilson et al., 2017]. Finally, excitatory neurons are more sparsely connected with each other, with a connection probability of around 10% [Harris and Shepherd, 2015], compared to inhibitory neurons, which target almost all neighboring excitatory neurons [Pfeffer et al., 2013].

It is not clear whether these differences across excitatory and inhibitory neurons serve computational purposes. It is also unknown whether these differences are all independent properties, each contributing to the computation, or if some differences are natural results of the others. To answer these questions, we first trained a deep recurrent convolutional neural network on an object classification task. Several structural principles of the brain are built into the network, including Dale’s law, the abundance of excitatory neurons, and the role of excitatory units as principal neurons. However, other observed properties of biological E-I circuits are not hardwired, and therefore could only evolve under the pressure of performing the task. We found that functional and structural differences across E and I neurons emerged through training. The development of these characteristics allows us to address the “why” question by removing the built-in differences across E and I neurons and monitoring whether specific functional and structural properties still emerge.

## 1 Multi-cell Convolutional Recurrent Network

The networks we use to model the visual cortex consist of 2 layers of purely feedforward, convolutional processing, followed by 2 layers of recurrent processing (Figure 1). The feedforward layers correspond loosely to retina and thalamus, while the recurrent layers correspond to cortex. Each recurrent layer consists of excitatory and inhibitory neurons (channels).

**Figure 1:**
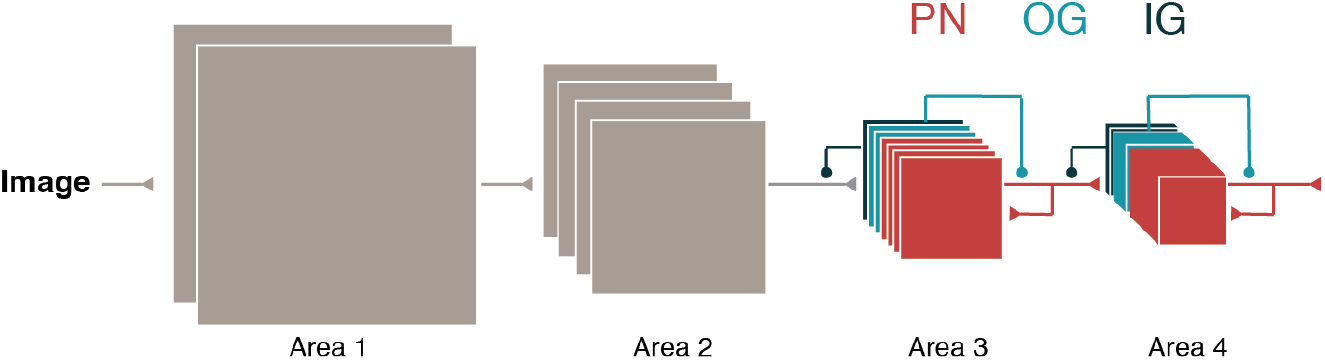
Convolutional recurrent network with multiple cell types. The neural network starts with two layers of regular convolutional processing, followed by 2 layers of recurrent processing. Each recurrent layer consists of three types of cells. In our standard implementation, the excitatory principal neurons (PN) output non-negative connection weights and target the next area. Output-gating (OG) and input-gating (IG) neurons are inhibitory and only target neurons within the same area.

In the brain, excitatory and inhibitory neurons can be further divided into many subtypes that differ in their inputs, output targets, and gene expression [Markram et al., 2004]. Two major types of inhibitory neurons account for 60% of all inhibitory neurons in the cortex [Rudy et al., 2011]. The first type expresses the molecule Somatostatin (SST) and inhibits dendrites of excitatory neurons, the structure that receives inputs from other neurons. The second type expresses the protein Parvalbumin (PV) and inhibits soma of excitatory neurons, where outputs are generated. Experimental evidence suggested that these input- and output-controlling inhibition can function like subtractive or multiplicative gates [Lee et al., 2012, Wilson et al., 2012]. As was previously pointed out [Costa et al., 2017], this motif in which principal neurons–those that can project to other areas–are recurrently controlled by multiple gates is reminiscent of the design of common recurrent units in machine learning, including Long Short-Term Memory (LSTM) [Hochreiter and Schmidhuber, 1997] and Gated Recurrent Unit (GRU) [Cho et al., 2014] networks.

Here we take this analogy further by modifying the structure of a convolutional LSTM unit [Xingjian et al., 2015] to introduce two distinct types of inhibitory neurons into the network (Figure 1). In the original LSTM network, all gate variables (forget, input, and output gates) are generated recurrently from the principal neurons through a single-layer perceptron. Instead, we use multi-layer perceptrons (MLPs) with a single hidden layer to recurrently generate the input and output gates. The two additional sets of neurons in the hidden layers become the input-gating (IG) and output-gating (OG) neurons.

Taken together, the multi-cell convolutional recurrent network we used is described by

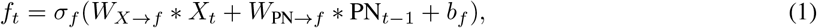

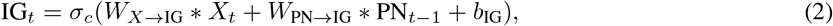

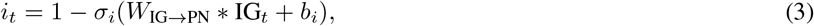

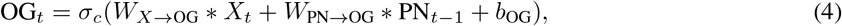

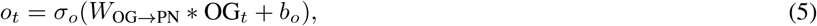

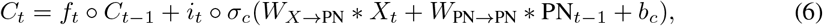

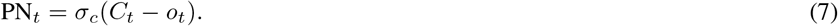

Here *f*_*t*_, *i*_*t*_, *o*_*t*_ stand for the forget, input, and output gates respectively, similar to LSTM. IG_*t*_, OG_*t*_, PN_*t*_ are the activity of the input-gating, output-gating, and principal neurons. *C*_*t*_ corresponds to the cell state in LSTM, and can be thought of as the pre-activation values (or membrane potential) of the principal neurons. All neurons are rectified linear units (ReLU), *σ*_*c*_ = max(*x*, 0). The activation function of the gate variable is the sigmoid function for multiplicative and forget gates (*σ*_*i*_, *σ*_*f*_), and ReLU for subtractive gates (*σ*_*o*_). *stands for convolution, while ⚬ stands for element-wise multiplication.

In the standard version of this network, we impose Dale’s law in the recurrent layers by ensuring that principal neurons (PN) only make excitatory connections – namely, all of their output weights are non-negative. This means the long-range connections from one recurrent layer to the next are also excitatory. In comparison, the gate neurons (IG, OG) all make inhibitory connections. Dale’s law is implemented by setting all relevant weight matrices *W*_PN*→•*_, *W*_IG*→•*_, *W*_OG*→•*_ to be non-negative using an absolute function, 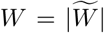, where the trainable variable is the non-sign-constrained weight matrix 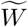. The connections for long-range inputs *W*_*X*→._ are positive if *X* originated from excitatory neurons. We found that task performance drops substantially if a rectified linear function is used to impose the sign constraint instead.

## 2 Reproducing functional and structural differences across cell types

We trained the multi-cell network on the image classification dataset CIFAR10 [Krizhevsky and Hinton, 2009]. We used common training techniques including momentum [Polyak, 1964] with an initial learning rate of 0.1 that decayed 10-fold at epochs 100, 150, and 200, as well as batch-normalization applied to the cell state *C*_*t*_. The network consists of two convolutional feedforward layers of 16, and 32 channels each, followed by two recurrent layers. The first recurrent layer contains 64 PN, 16 IG, and 16 OG channels. The second recurrent layer contains 128 PN, 32 IG, and 32 OG channels. The network is unfolded for 4 time steps, and the classification output is read-out from the final time step using a fully connected linear layer. See Appendix for more details. For all conditions, we trained 5 networks with different random seeds. The network reaches approximately 85% test accuracy on CIFAR10, comparable to convolutional networks of similar depth [Hinton et al., 2012].

We ask whether training develops qualitative differences between excitatory and inhibitory neurons in the network, as observed in the brain. In mouse cortex, inhibitory neurons are functionally less selective than excitatory neurons [Kerlin et al., 2010, Znamenskiy et al., 2018], meaning that they tend to respond to different stimuli with similar values. We first measured the selectivity to static oriented gratings for excitatory and inhibitory neurons (see Appendix for details). The selectivity is quantified in two ways. First, we computed the global Orientation Selectivity Index (gOSI) for each neuron *j*,

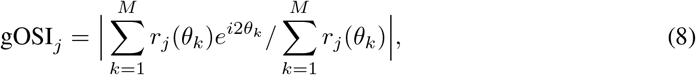

where *r*_*j*_(*θ*_*k*_) is the *j*-th neuron’s response to the *k*-th grating stimulus at orientation *θ*_*k*_. gOSI is close to 1 when the neuron is highly selective to orientation. For each type of neurons, we report the average gOSIs across all channels in the center of the convolutional layer. The second measure of selectivity is the skewness of the distribution of responses to various stimuli [Samonds et al., 2014]. The skewness *γ*_*j*_ for neuron *j* is defined as

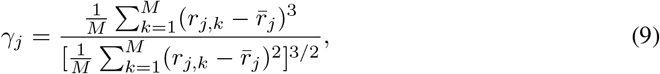

where *r*_*j,k*_ is the *j*-th neuron’s response to the *k*-th stimulus, and 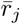 is the average response. The skewness is higher if a neuron is strongly selective to a small number of stimuli. For both gOSI (Figure 2a,b) and orientation skewness (Figure 2c), the selectivity is substantially higher for excitatory neurons (PN) compared to inhibitory neurons (IG, OG), in both recurrent layers (areas 3 and 4). Next, we measured the selectivity to natural images. We computed the skewness of responses to all images in the test set. Again, excitatory neurons in both areas 3 and 4 have higher selectivity to natural images than inhibitory neurons (Figure 3).

**Figure 2:**
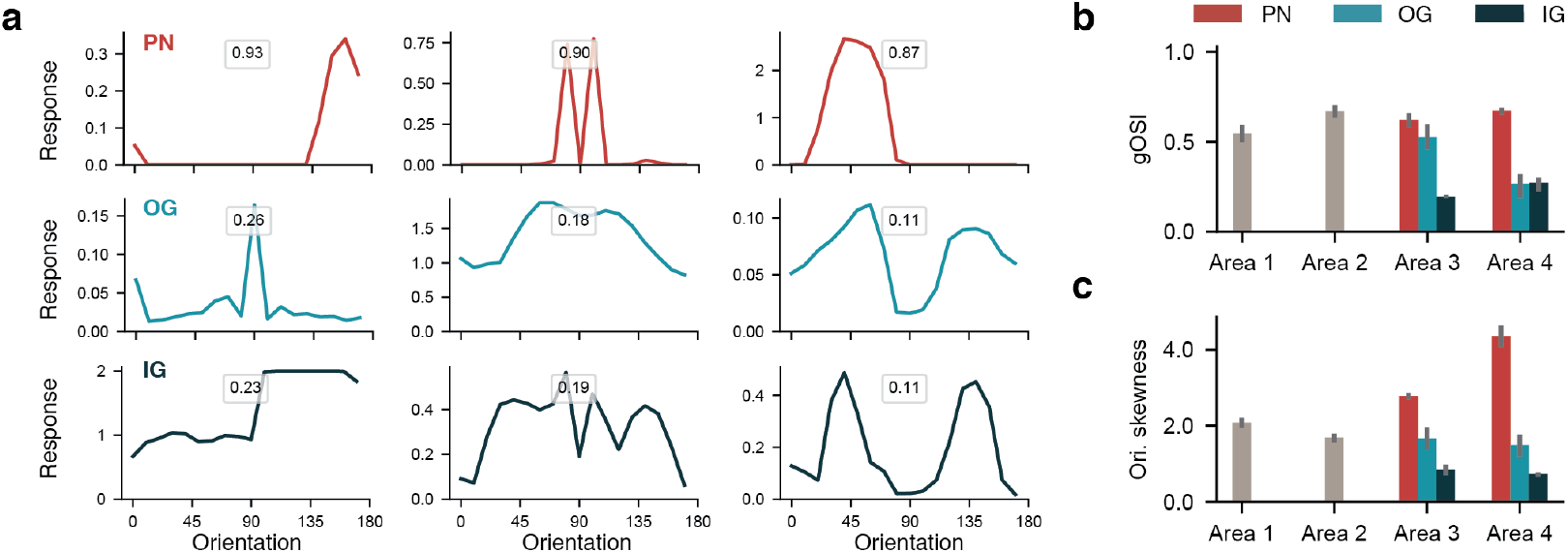
Model excitatory neurons have higher orientation selectivity than inhibitory neurons. (a) The orientation tuning curves of example PN, OG, and IG neurons from area 3 of one standard network. The number in each panel indicates the neuron’s global Orientation Selectivity Index (gOSI). The standard model is highlighted in bold. (b, c) The average gOSI (b) and orientation skewness (c) for each type of neurons in the standard networks. Error bar is the 95% confidence interval computed with bootstrapping across 5 networks (same for following figures).

**Figure 3:**
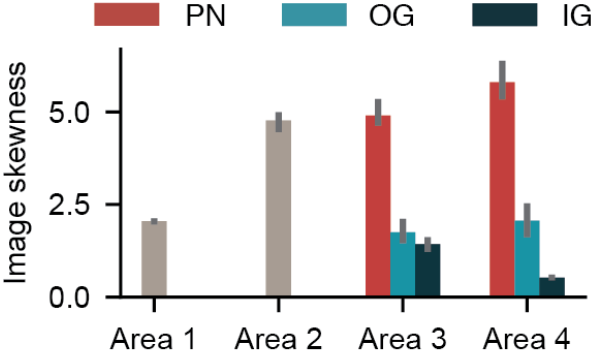
Excitatory neurons have higher natural image selectivity. The selectivity is measured with the skewness of the distribution of responses to natural images.

Another major qualitative difference between excitatory and inhibitory neurons in the brain is the higher density of I-to-E connections compared to E-to-E. In our network, we found that the distribution of all recurrent connection weights is multi-modal (Figure 4a), with a group of strong connections and many much weaker connections. To assess the connection density, we chose a threshold at exp(−10), and quantified the proportion of connection weights that exceed this threshold. This threshold is chosen to separate the strongest mode from the weaker modes in the distribution of all weights (Figure 4a). The distribution of PN-to-PN connections is spread out (Figure 4b), leaving a substantial proportion of connections below threshold. In contrast, the distributions of both IG-PN and OG-PN connection weights are mostly concentrated above the threshold (Figure 4c,d), leading to higher connection density (Figure 4e). Therefore, sparser excitatory connectivity emerged in the network after training, in agreement with biological observations.

**Figure 4:**
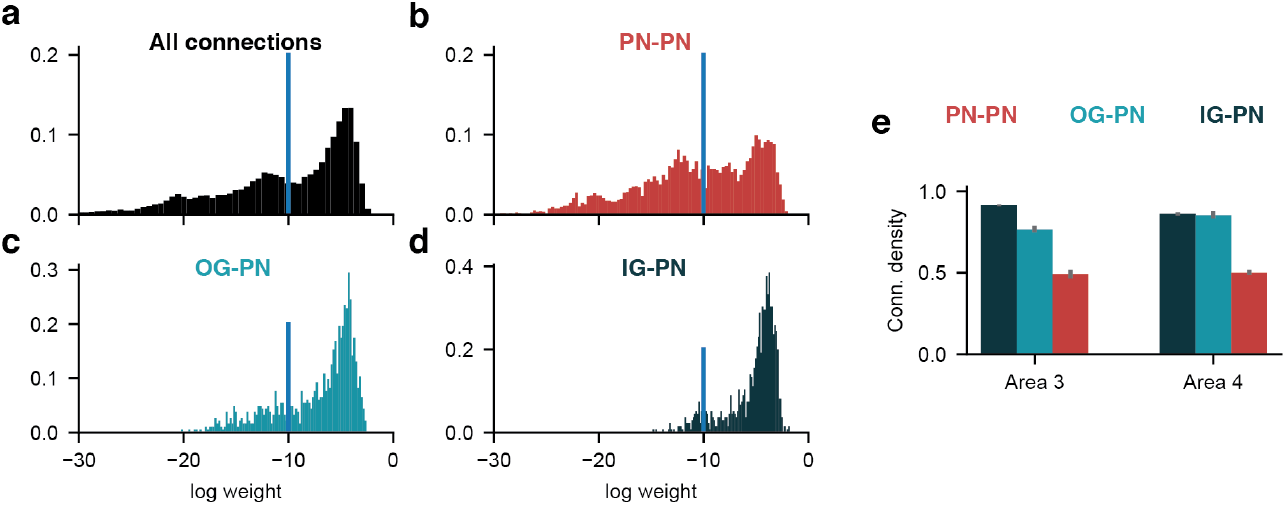
Model excitatory neurons are more sparsely connected. (a-d) The distribution of connection weights for all recurrent connections (a), and for the PN-to-PN (b), OG-to-PN (c), and IG-PN connections (d) in area 3 of one standard network. Blue line: the threshold used to compute the proportion of strong connections, namely those that exceed the threshold. (e) The connection density (proportion of strong connections) for three types of excitatory and inhibitory connections across area 3 and 4.

To test the generality of these results, we trained networks with subtractive or multiplicative gates (Figure 5a), networks with only one type of inhibitory neurons (Figure 5b), and networks with or without batch normalization or dropout on the three cell types (Figure 5c). In all variations of networks that reached accuracy above 80%, the excitatory neurons are more selective and less connected. Preliminary results from networks trained on ImageNet [Deng et al., 2009] showed the same qualitative findings.

**Figure 5:**
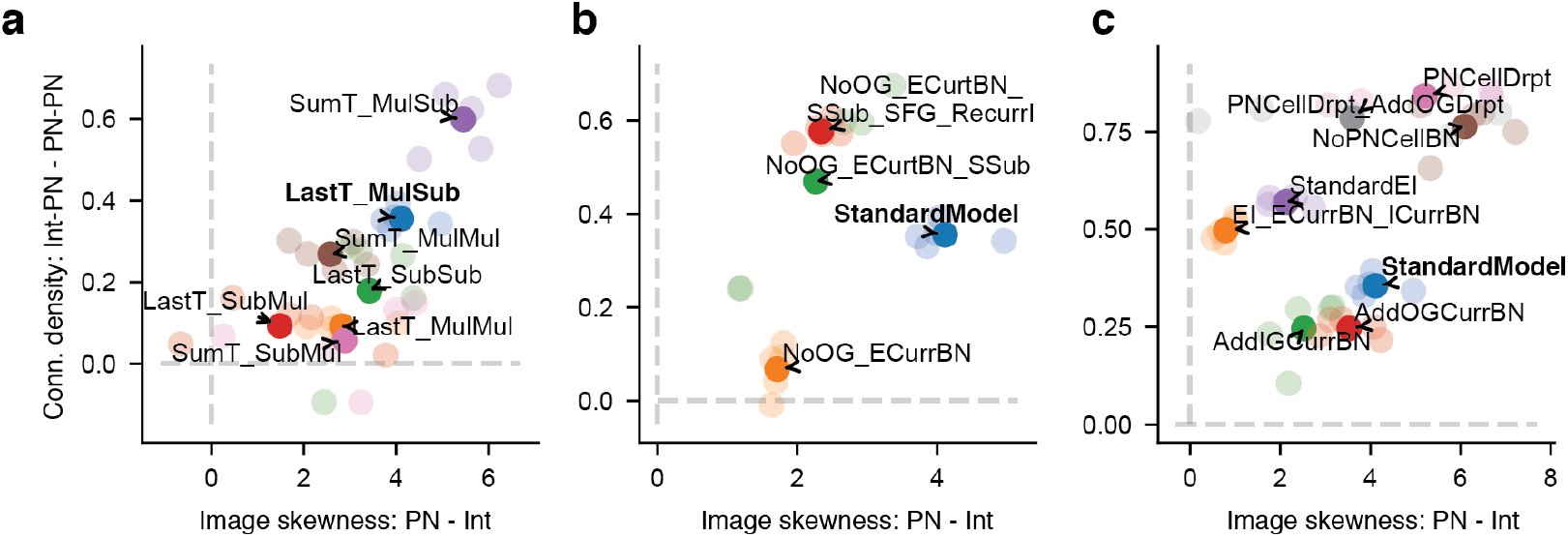
The E-I differences emerge across a range of network variations. (a-c) The E-I differences are summarized using the difference in image skewness between principal neurons (PN) and interneurons (Int), and the difference in connection density between interneuron-to-PN and PN-to-PN connections. (a) Models with different readout and gating mechanisms. (b) Models in transition from the standard network to a more simplified E-I network. (c) Models with variations of batch normalization or dropout (c) all develop the E-I differences. Results are combined from areas 3 and 4. Light circles: individual networks. Dark circles: average. See Appendix for details.

By monitoring the orientation selectivity, image selectivity, and connection density throughout the training process, we found that the differences across excitatory and inhibitory cells emerged early on (Figure 6). Even though excitatory and inhibitory neurons started out with similar selectivity (close to 0), and similar connectivity (close to 1), the differences across cell types become substantial after 5-30 epochs. These functional and structural differences remain stable throughout training, even as the training performance continues to improve (see Appendix). Together, these results argue that there exists a strong optimization pressure to differentiate the selectivity and connectivity of excitatory and inhibitory neurons.

**Figure 6:**
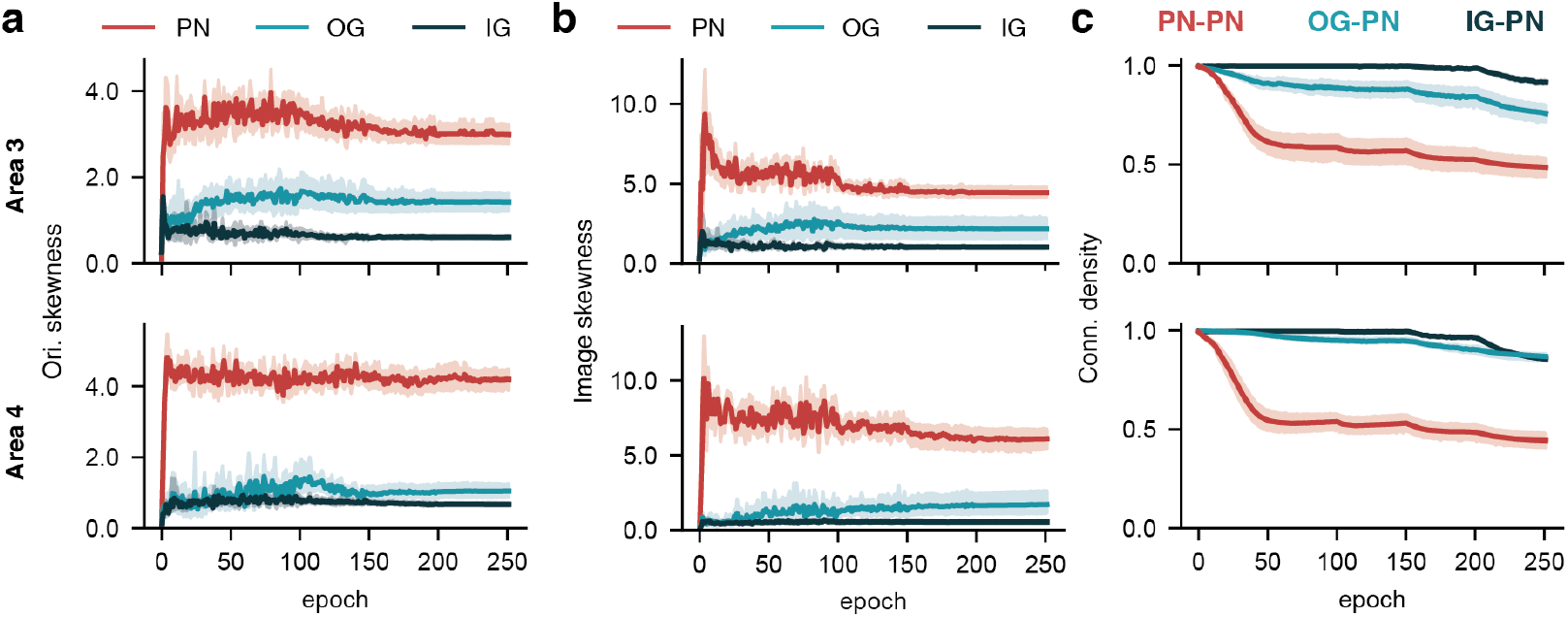
The E-I differences emerge early in training. The orientation skewness (a), image skewness (b), and connection density (c) for areas 3 (top) and 4 (bottom) throughout training.

## 3 Why do excitatory and inhibitory neurons have distinct selectivity and connectivity?

We have shown that neural networks with excitatory and inhibitory neurons develop different selectivity and connectivity, qualitatively reproducing long-standing findings in the brain. Now we ask why these differences emerge.

The emergent differences across our model E and I neurons can only be explained by their built-in structural asymmetry. We have incorporated three major forms of asymmetry that exist in the brain. First, there is an asymmetry in numbers. Excitatory neurons are 4 to 10 times more abundant than inhibitory neurons in the brain. In the standard network, there are 4 times more PNs compared to IGs or OGs in each area. Second, there is an asymmetry in projection targets. In the cortex, all principal neurons are excitatory. Meanwhile, all inhibitory neurons are interneurons, meaning that they only connect to other neurons within the same area. Third, there is by definition an asymmetry in action. Excitatory neurons excite other neurons while inhibitory neurons inhibit. When the activation function of a neuron is rectified and non-saturating (for example, ReLU), excitatory inputs can move the neuron into a responsive regime, where inhibitory inputs can make a neuron non-responsive. In this section, we will remove individual asymmetry, and test which one led to the observed differences in selectivity and connectivity. We will present results based on variations of the standard model, but all results are qualitatively reproduced in a simplified E-I network (see Appendix).

### Asymmetry in numbers

In our standard network, the ratio between the number of OGs(IGs) and PNs is 1:4. It is conceivable that excitatory neurons can afford to be more selective to orientations and images because there are more of them. To test this hypothesis, we varied the number of inhibitory channels in the recurrent layers, while maintaining the number of excitatory channels. The OG/IG:PN ratio tested ranges from 1:64 to 4:1. We found little evidence that the difference in E and I selectivity depends on the ratio of their numbers (Figure 7). The orientation and image selectivity are still higher among excitatory neurons even when there are 4x more inhibitory neurons of each type. In comparison, the density of inhibitory connections decreases as the number of inhibitory channels increase, while the connection density of excitatory neurons remain steady.

**Figure 7:**
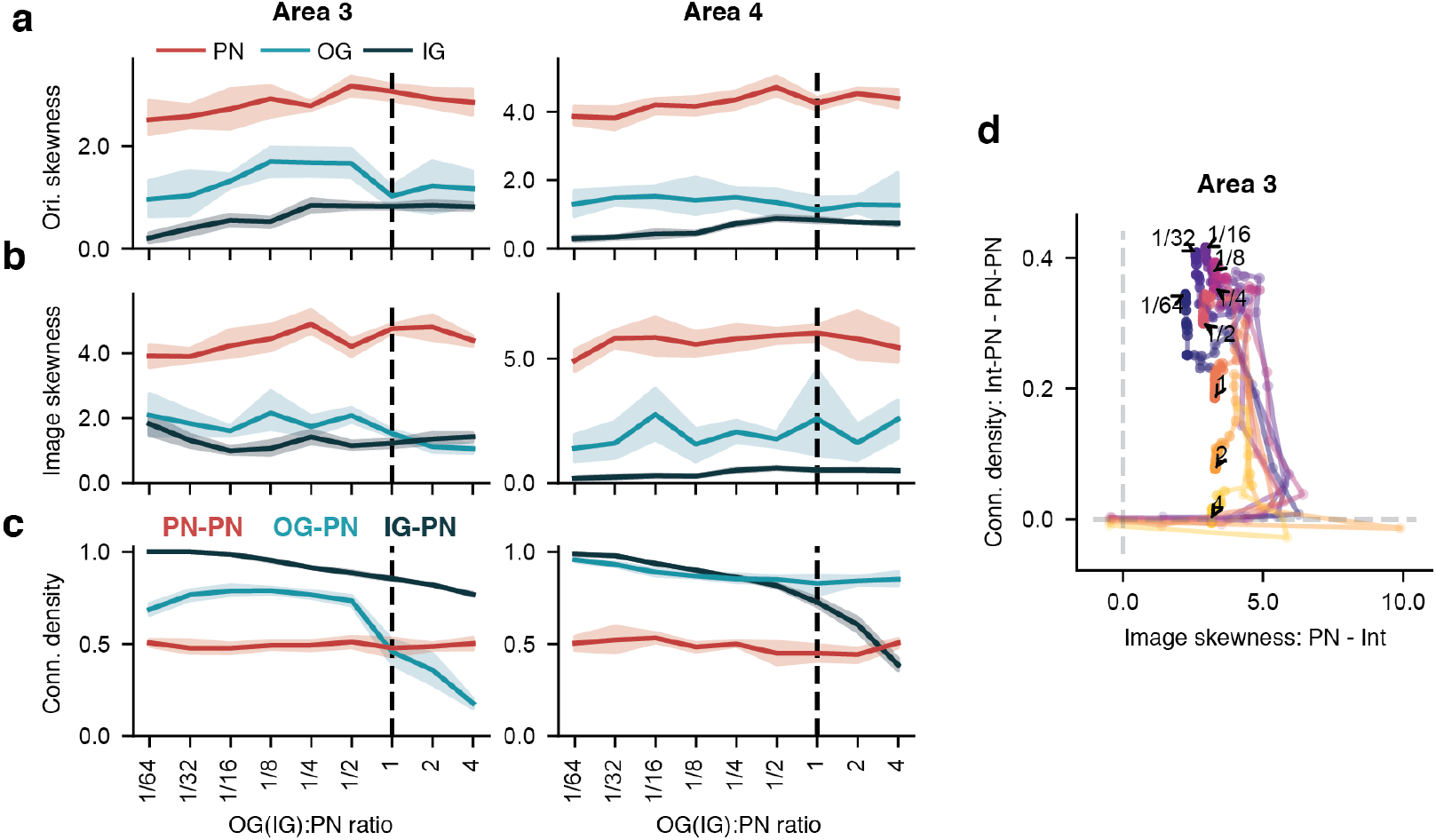
Selectivity and connectivity in networks with various ratios of excitatory and inhibitory neurons. (a-c) The orientation skewness (a), image skewness (b), and connection density (c) for networks where the number of inhibitory neurons (both IG and OG) varied from 1/64 to 4x of the number of excitatory neurons. (d) The E-I difference in connection density versus the difference in image skewness when the OG/IG:PN ratio is varied. Each colored curve shows how networks evolved in this E-I difference space through training. All networks started around the origin (no E-I differences). Difference in selectivity evolved first, followed by difference in connectivity.

### Asymmetry in projection

In both the cortex and our standard network, principal neurons are exclusively excitatory. Here, we decoupled this relationship by training *InhPN* networks with inhibitory principal neurons and excitatory interneurons. The connections from principal neurons to the next layer are kept excitatory in these networks, because we had difficulty training networks with inhibitory long-range projections past 80% accuracy. InhPN networks can still achieve accuracy similar to that of the standard network (Figure 8a). However, the excitatory neurons (interneurons) are less selective to orientation and natural images compared to the inhibitory neurons (principal neurons) (Figure 8b,c, Appendix). This result argues that higher selectivity is not a property inherent to excitatory neurons. Whichever type of neuron serves as the principal neurons would demand higher selectivity, presumably because the principal neurons need to carry detailed stimulus information to the next layer.

**Figure 8:**
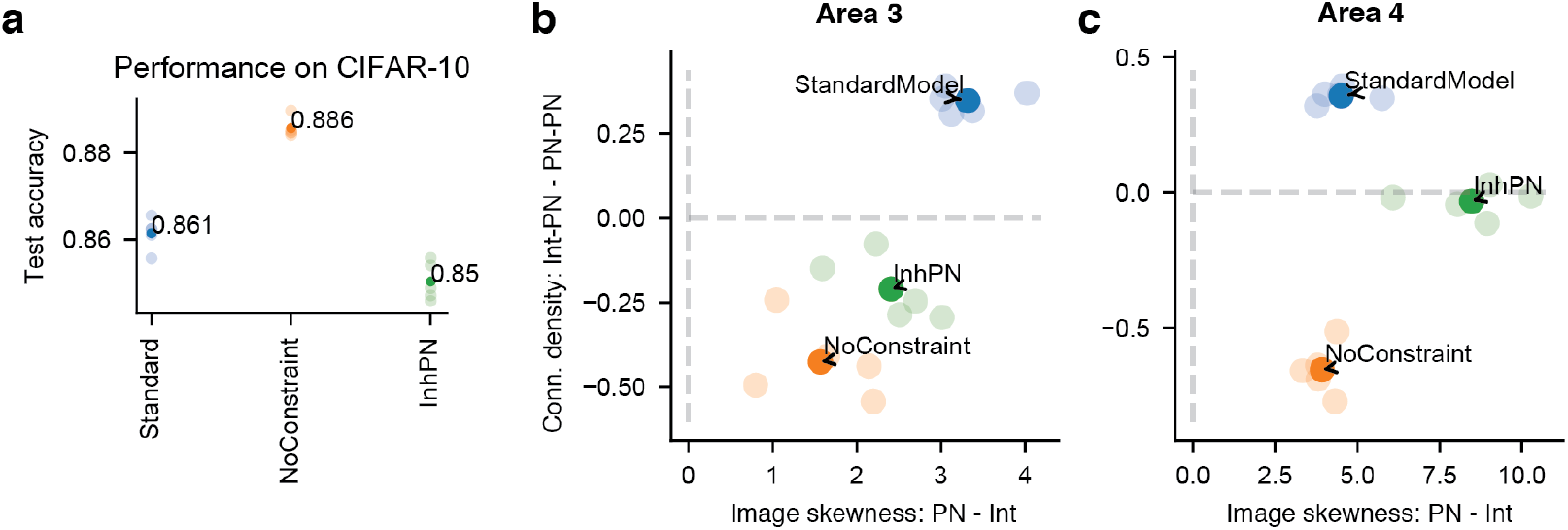
Removing asymmetry between excitatory and inhibitory neurons. (a) The performance of 3 types of networks on CIFAR10. (b,c) Difference in connection density against difference in image selectivity for areas 3 (b) and 4 (c) of three types of networks. Light circles: individual networks; dark circles: average.

Interestingly, in the InhPN networks, the connectivity among inhibitory (principal) neurons remains dense despite a heightened selectivity (Figure 8b,c, Appendix). This result is in stark contrast with the sparser connectivity needed for highly selective excitatory neurons in the standard network. These seemingly contradictory results can be understood in the context of recurrently connected ReLU neurons. When a ReLU neuron is highly selective, typically, it is strongly activated by only a small subset of stimuli. If this selectivity is supported by recurrent excitatory connections, then a neuron should receive inputs mainly from a small set of excitatory neurons with similar preferred stimuli. Meanwhile, if the selectivity is supported by recurrent inhibition, then a neuron should receive inputs from nearly all other neurons, except for the small subset of neurons with similar preferences. Therefore, recurrent connections between highly selective inhibitory neurons should be dense, instead of sparse.

### Asymmetry in action

The namesake difference between excitatory and inhibitory neurons is the sign of their connection weight values. All connections stemming from an excitatory (inhibitory) neuron are constrained to be non-negative (non-positive). We can release this constraint on the sign of connection weights, while keeping other properties of the standard network. Such *NoConstraint* networks achieve slightly better accuracy compared to the standard Dale’s law-obeying networks (Figure 8a). In the NoConstraint networks, the formerly-excitatory principal neurons developed higher selectivity compared to the formerly-inhibitory interneurons (Figure 8b,c), consistent with our previous finding that principal neurons have higher selectivity regardless of the sign of their outputs.

Similar to previous results, in the NoConstraint network, the principal neurons no longer have sparser connectivity, instead, they are almost fully connected (Figure 8b,c, Appendix). This result again emphasizes that the link between connectivity and selectivity depends crucially on whether Dale’s law is applied. In a network without Dale’s law, a neuron can receive inputs from all other neurons and remain selective, as long as the inputs from most neurons cancel out, leaving this neuron effectively driven by a small set of inputs.

## 4 Discussion

We have shown that recurrent neural networks equipped with multiple cell types are capable of capturing several important features of the brain, including higher selectivity and sparser connectivity among excitatory neurons. These qualitative features emerged solely from the pressure to perform the task, suggesting that these qualities are indeed beneficial to task performance. This allows us to study what aspects of the network give rise to this distinction between excitatory and inhibitory neurons. We found that the higher selectivity of excitatory neurons is mainly driven by their role as the principal neurons that transmit information to upper layers. When Dale’s law is obeyed, a higher selectivity necessitates sparser connectivity among excitatory neurons.

Optimization algorithms like stochastic gradient descent combined with large datasets are highly effective at tuning connection weights to perform tasks. However, it remains unlikely to observe a specific set of desired structural principles emerge naturally through training. We designed our standard network to obey Dale’s law. Theoretically, connectivity respecting Dale’s law could emerge out of training, but has not been demonstrated, to the best of our knowledge. There are at least three possibilities: (1) the optimization algorithm is not strong enough to discover this solution because it remains a exponentially small part of the solution space (1*/*2^*N*^ of the entire solution space, *N* being the number of connection weights). Similarly, it would be difficult for a ResNet structure [He et al., 2016] to develop from training a vanilla 100+ layer deep feedforward network. (2) The task we used is not appropriate for the emergence of Dale’s law. It is conceivable that Dale’s law is beneficial to some tasks that have yet been identified. (3) Finally, it is possible that Dale’s law is a result of a compromise that the brain has to strike due to its biological nature, and is irrelevant to general computing machines. Understanding the nature of the computational benefit of Dale’s law (if any) would be a major achievement in computational neuroscience, and may shed light on better designs of artificial neural networks.

