## Appendix for "Understanding the functional and structural differences across excitatory and inhibitory neurons"

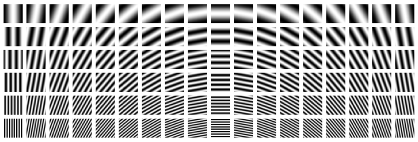

Figure S1: **The static gratings.** A total of 432 gratings, using 18 orientations (10° apart), 6 spatial frequencies (1, 2, 3, 4, 6, 8) and 4 phases (90° apart).

### A Additional details of the models

Our models consist of two feedforward layers and two recurrent layers. Convolutions in the feedforward layers are regular convolutions. In the recurrent layers, we use depth-separable convolution, where a spatial convolution is applied on each channel separately and then followed by a point-wise convolution that mixes channels [Chollet, 2017]. Kernel size is 3x3 for all convolutions. To measure the connection density, the 4-d convolution matrix of size [3, 3, channels_in, channels_out] is recovered from the spatial and point-wise convolution kernels, and then averaged over the first two spatial dimensions. This would give us a channel-to-channel weight matrix, with a total of channels_in×channels_out connection weights. The results are similar when we do not average across space and instead analyze the 3×3×channels_in×channels_out connection weights. Density is quantified as the proportion of connection weights that exceed a chosen threshold. We picked a threshold $\exp(-10)$ that separates the strong mode from the weaker modes in the distribution of all connection weights. The precise value of the threshold does not impact our qualitative results. Max pooling is applied on the output of each layer. Pooling stride is 1 for the first feedforward layer, and 2 for other layers.

### B Computing orientation selectivity

A static oriented grating is a two-dimensional sinusoidal wave $G(x, y)$ satisfying:

$$G(x, y) = \cos\{k[-(x - x_0)\sin\theta + (y - y_0)\cos\theta] + \phi\},$$

where $\theta$ is the orientation, $k$ is the spatial frequency, $\phi$ is the phase and $(x_0, y_0)$ is the center location. We generate a total of 432 gratings (Figure S1) using 18 orientations (10° apart), 6 spatial frequencies (1, 2, 3, 4, 6, 8) and 4 phases (90° apart).

We present each grating image to a network, and each neuron's preferred orientation, spatial frequency and phase is chosen when the neuron has the maximal activity. At its preferred spatial frequency and preferred phase, the global Orientation Selectivity Index (gOSI) is computed based on all orientations. Unlike gOSI, the orientation skewness is computed based on all 432 grating images.

### C Model variants

Here we describe the model variants trained for Figure 5.

#### C.1 Readout and gating

In our standard model, the logits are read out from the last time step of the final recurrent layer (area 4). We also trained networks where the logits are read out from the final recurrent layer's activity summed across all time steps. In Figure 5a, We denote the former structure as "LastT", and the latter structure as "SumT".

Apart from multiplicative (Mul) input gates and subtractive (Sub) output gates in the standard network, we implemented other combinations of gating mechanisms. In Figure 5a, a network is named as "Readout_InputGateOutputGate". For example, our standard model is named "LastT_MulSub" in this plot.

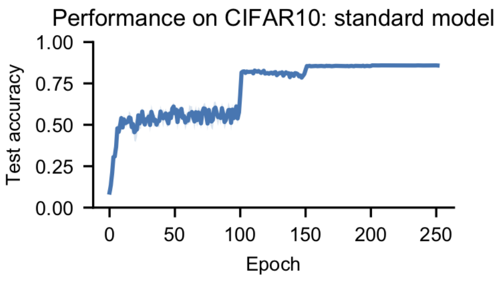


Figure S2: **The classification accuracy during training.** The learning rate is decayed at epoch 100, 150, and 200.

##### C.2 E-I model

We refer to models with only one type of inhibitory neurons as E-I models. Our standard E-I model is implemented as:

$$\begin{aligned}
\mathrm{Curr}_E &= W_{F\to E} * X_t + W_{E\to E} * E_{t-1} - W_{I\to E} * I_{t-1} + b_E \\
\mathrm{Curr}_I &= W_{F\to I} * X_t + W_{E\to I} * E_{t-1} - W_{I\to I} * I_{t-1} + b_I \\
E_t &= f_E \circ E_{t-1} + (1 - f_E) \circ \sigma_c(\mathrm{Curr}_E) \\
I_t &= f_I \circ I_{t-1} + (1 - f_I) \circ \sigma_c(\mathrm{Curr}_I)
\end{aligned}$$

$E_t$ and $I_t$ are the activity of the excitatory and inhibitory neurons. $\mathrm{Curr}_E$ and $\mathrm{Curr}_I$ are the input currents for E and I neurons respectively, with batch normalization applied to $\mathrm{Curr}_E$. $f_E$ and $f_I$ are forget gates, implemented as $\sigma(\widetilde{f})$ where $\sigma$ is the sigmoid function and $\widetilde{f}$ is a trainable 1-d tensor with the same size as the number of channels. Excitatory neurons are also principal neurons in this model.

Figure 5b contains a set of models in transition from our standard model to this E-I model. *NoOG_ECurrBN* removes the output gate of our standard model and moves the batch normalization from cell state $C_t$ to the input current of PN neurons. Based on *NoOG_ECurrBN*, we obtained *NoOG_ECurrBN_SSub* by changing the multiplicative input gate into a simplified subtractive current. By furthering simplifying the forget gate structure to be the same as the E-I model, and adding recurrent structure for IG neuron, we get the *NoOG_ECurrBN_SSub_SFG_RecurrI* model, which is equivalent to the standard E-I model.

##### C.3 Batch normalization and dropout

In Figure 5c, we tested various networks with or without batch normalization or dropout. In our standard model (*StandardModel*), removing the batch normalization on cell states (*NoPNCellBN*) or replacing it with a dropout function (*PNCellDrpt*) (keep probability equals to 0.9) retains the performance above 80% and the E-I differences in both selectivity and density (Figure 5c).

For the standard model with gating neurons, adding batch normalization for IG (*AddIGCurrBN*) or OG neurons (*AddOGCurrBN*) increases selectivity for the batch-normalized inhibitory neurons. Nevertheless, the inhibitory selectivity remains lower than that of PNs. The same trend is observed when batch normalization is added to inhibitory neurons in the standard E-I model (*EI_CurrBN_ICurrBN*), or when dropout is added to output-gating neurons in *PNCellDrpt* (*PNCellDrpt_AddOGDrpt*).

#### D Classification performance during training

The classification performance of the standard network increases rapidly in the first 10 epochs, and continues to improve after 150 epochs of training. The E-I difference in selectivity also increases rapidly in the first 10 epochs (Figure 6a,b), while the difference in connection density plateaus at about epoch 50 (Figure 6c).

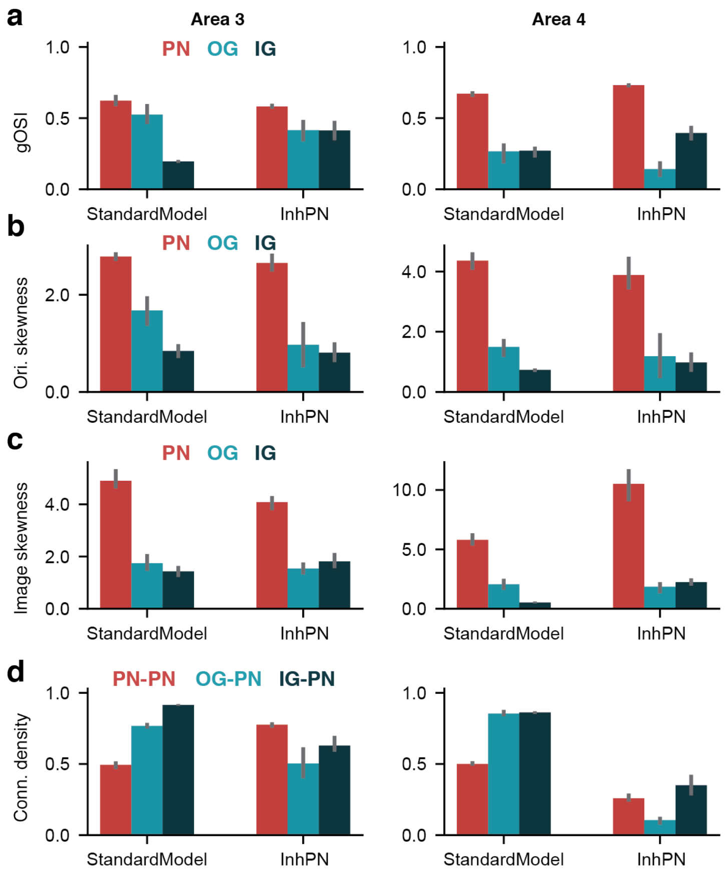


Figure S3: **Detailed comparison between standard model and InhPN network.** Comparing gOSI (a), orientation skewness (b), image skewness (c), and connection density (d) for areas 3 (left) and 4 (right) between standard model and the InhPN network.

#### E Detailed comparison between the standard model and its variants

The standard model is compared to the InhPN network (Figures S3) and the NoConstraint network (Figure S4).

#### F Analysis of the standard simplified E-I model

We trained simplified networks with a single type of inhibitory neurons, as described above. The standard E-I model shows similar qualitative trends (Figure S5) as the standard network used in the main text. InhPN and NoConstraint variants of the standard E-I model behave similarly to InhPN and NoConstraint versions of the standard network (Figure S6).

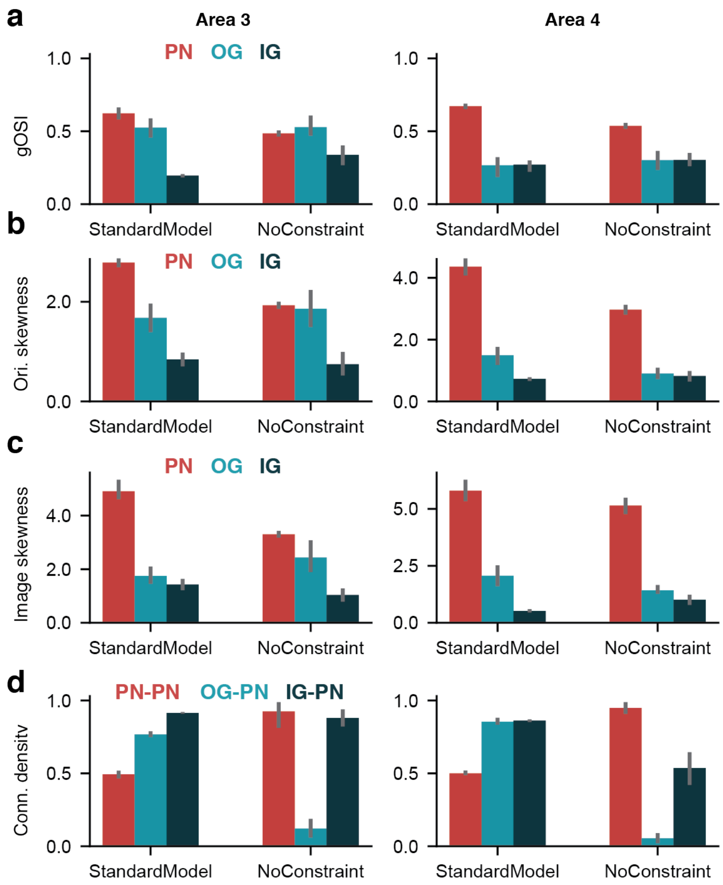


Figure S4: **Detailed comparison between standard model and NoConstraint network.** Comparing gOSI (a), orientation skewness (b), image skewness (c), and connection density (d) for areas 3 (left) and 4 (right) between standard model and the NoConstraint network. In the NoConstraint model, the OG-PN connectivity is close to zero, suggesting that OGs are only weakly participating in the computation.

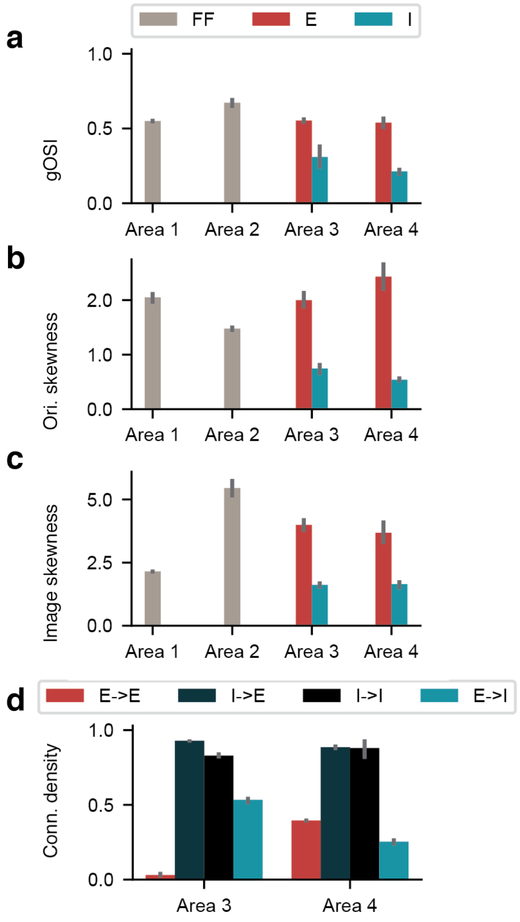


Figure S5: **Standard E-I model.**

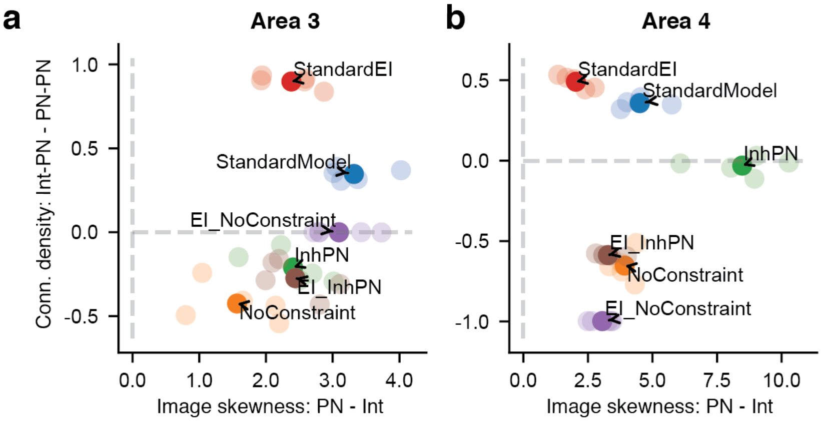


Figure S6: **InhPN and NoConstraint variants of the standard E-I model.**
